# Mosquito densoviruses: the revival of a biological control agent against urban *Aedes* vectors of arboviruses

**DOI:** 10.1101/2020.04.23.055830

**Authors:** Aurélie Perrin, Anne-Sophie Gosselin-Grenet, Marie Rossignol, Carole Ginibre, Bethsabée Scheid, Christophe Lagneau, Fabrice Chandre, Thierry Baldet, Mylène Ogliastro, Jérémy Bouyer

## Abstract

Urban *Aedes* mosquitoes are vectors of many viruses affecting human health such as Dengue, Chikungunya and Zika viruses. Insecticide resistance and environmental toxicity risks hamper the effectiveness of chemical control against these mosquito vectors. Alternative control methods, such as the use of mosquito-specific entomopathogenic viruses should be explored. Numerous studies have focused on evaluating the potential of different densoviruses species as biological control agent. However, knowledge on the extent of inter- and intra-specific variations in the susceptibility of *Aedes* mosquitoes to infection by different densoviruses remains insufficient. In this study, we compared infection and mortality rates induced by the Aedes albopictus densovirus 2 in different strains of *Aedes albopictus* and *Aedes aegypti* mosquitoes. The two *Aedes* species were different in terms of susceptibility to viral infection. Under laboratory conditions, Aedes albopictus densovirus 2 appeared more virulent for the different strains of *Aedes aegypti* tested than for those of *Aedes albopictus*. In addition, we also found significant intra-specific variation in infection and mortality rates. Thus, although even if Aedes albopictus densoviruses could be powerful biocontrol agents used in the management of urban *Aedes* populations, our results also call into question the use of single viral isolate as biocontrol agents.

## Introduction

*Aedes albopictus* (the tiger mosquito) and *Ae. aegypti* (the yellow fever mosquito), are particularly invasive species that proliferate in tropical and temperate urban environments and are the main vectors of Dengue, Chikungunya, Yellow fever and more recently Zika viruses. In the context of globalization and the movement of goods and people, these emerging vector-borne diseases are now present on almost every continent^1,2^. In the absence of vaccine or antiviral therapy for the majority of these diseases, vector control is the main strategy to prevent their spread. This is mainly practiced by controlling adult mosquito populations through spatial treatments using pyrethroid-based chemical insecticides and by controlling larvae through physical suppression of breeding sites or larvicides. The application of insecticides can be problematic because of their high environmental and human health toxicity^3–6^, their general toxicity to non-target insects^7^, and the insecticide resistance of target mosquitoes^8,9^. Pyrethroids are the most widely used chemical insecticides in the world but their intensive use has led to the selection of pyrethroid resistant mosquitoes worldwide^10–12^. Many innovative approaches are being developed to control *Aedes* sp. mosquitoes such as adult traps, lethal ovitraps, autodissemination stations, Release of insects with dominant lethality (Ridl), Sterile Insect Technique, Incompatible Insect Technique^13^ but larval control remains essential and is systematically includes in any integrated control strategy. The control of urban *Aedes* larvae is extremely complex to implement because of the diversity and multitude of larval habitats, which are made up of small, and usually cryptic, water containers^14^. Apart from chemical larvicides (e.g. temephos, pyriproxyfen, diflubenzuron), the biological larvicide recommended against urban *Aedes* larvae is derived from *Bacillus thuringiensis* subsp. *israelensis* (Bti), a natural soil bacteria selected for its exclusive pathogenic action on some species of Diptera^15^. However, its effectiveness is limited by many biological and environmental factors: sunlight, amount of organic matter, larval density and depth of breeding sites^16^. For the moment, no resistance to Bti has been observed in mosquito but it is crucial to develop other candidate larvicides to fulfil the range of effective and environmental-friendly control tools. The use of several bio-larvicides with different action spectra should ensure effective, feasible and sustainable vector control and should contribute to manage the resistance of target insects to the active molecules.

Many viruses are known to be pathogenic for mosquitoes^17–19^, but their potential use in biological control has been limited by their low infectivity or a production method unsuitable for field treatment. Mosquito densoviruses (MDVs), that exhibit a narrow host range and multiple transmission patterns, are, however, a potential alternative^20,21^. Densoviruses (DVs), also known as densonucleosis viruses, are small icosahedral non-enveloped DNA viruses belonging to the *Parvoviridae* family and are highly infectious for invertebrate (insects, crustaceans and echinoderm)^22,23^. They are divided into 5 genus^24^: *Ambidensovirus, Brevidensovirus, Iteradensovirus, Hepandensovirus* and *Penstyldensovirus*. MDVs have been isolated from laboratory colonies or natural populations of mosquitoes and from chronically infected mosquitoes-derived cell lines. They have been classified in the first two genus. Within the *Brevidensovirus* genus, there are currently two type species with 9 virus species i) the *Dipteran brevidensovirus 1* with Aedes aegypti densovirus 1 and 2 (AaeDV1^25^ and AaeDV2^26^ respectively), Aedes albopictus densovirus 1 (AalDV1^27^), Culex pipiens pallens densovirus (CppDV^28^) and Anopheles gambiae densovirus (AgDV^29^), ii) the *Dipteran brevidensovirus 2* with Aedes albopictus densovirus 2 and 3 (AalDV2^30^ and AalDV3^31^ respectively) and Haemagogus equinus densovirus (HeDV^32^). In the genus *Ambidensovirus*, only one strain has been described, the Culex pipiens densovirus (CpDV^33^). Five others viruses are described in literature but are not yet included in the official taxonomy. Three strains of Aedes albopictus densoviruses, AalDV4^34^ to AalDV6 have been isolated and described from C6/36 cell line without any cytopathic effect suggesting that persistent cryptic infections are common^32^. The sequences of these viruses are sufficiently different that it is highly unlikely that they have evolved from a single contamination event^35^. Despite the lack of cytopathic effect in cell lines, theses MDVs have been shown to be pathogenic to mosquito larvae by oral infection and are able to replicate and to be transmitted in adult mosquitoes. Unlike strains isolated from cell lines, a new strain, AalDV7, was isolated from field-collected *Ae. albopictus*^36^. Actually, the Aedes Thai strain densovirus (AthDV) was detected in colonies of *Ae. aegypti* and *Ae. albopictus* from Thailand^37^. MDVs can infect a wide range of mosquitoes, but natural infection appears to be confined to a single host species. The host range has been more or less well described according to the viruses. AaeDV1, the best characterized of them, was infectious in laboratory experiments for *Ae. aegypti, Ae. albopictus, Ae. cantans, Ae. caspius, Ae. geniculatus, Ae. vexans, Cx. pipiens* and *Culiseta annulata*^38^. MDVs are thought to persist in nature by horizontal transmission from larvae to larvae in the wild aquatic environments, although transovarial and sexual transmission have also been recorded^3,21,29,36,39–42^. They are highly pathogenic for larvae at all stages, but mortality is higher when infection occurs at an early stage. Older larvae can survive and grow into imago after the virus infection, and infected adult female mosquitoes can transmit the virus vertically to the next generation^43^. Mortality is also higher during critical phases of mosquito life, especially during larval metamorphosis, pupation and adult emergence, which require more energy.

MDVs are emerging as promising tools for the control of *Aedes* mosquito population. The objective of the ERC Revolinc project, that funded this study, is to use these biopesticides to boost the Sterile Insect Technique^44^. Sterile males would thus be coated with these viruses before being release, thus contaminating wild females even in the absence of successful mating^44^. Most studies have been devoted to evaluating the potential of different MDVs strains as biocontrol agents. Knowledge on the extent of inter- and intra-specific variations in the susceptibility of *Aedes* mosquitoes to DVs infection are lacking. The overall objective of this study was to determine the potential of AalDV2 as a biological control agent against *Aedes* urban mosquitoes. AalDV2 (formerly known as AaPV for Aedes albopictus parvovirus) was isolated from a chronically infected cell line of the C6/36 clone of *Ae. albopictus*^30,45,46^ and was found to be highly pathogenic for *Ae. aegypti* neonate larvae reaching 95% mortality in the N’ Goye strain. No data are available on the pathogenicity of this virus to *Ae. albopictus*. In this work, we assessed the pathogenicity of AalDV2 against different strains of *Ae. albopictus* and *Ae. aegypti* mosquitoes from different geographical areas.

## Results

### Susceptibility of different strains of urban *Aedes* to AalDV2 infection

#### A high level mortality in larvae and pupae

Three replicates of 150 first-instar larvae of each strain of *Ae. albopictus* and *Ae. aegypti* were, or not, exposed to AalDV2. Larval mortality appeared 5 to 7 days after infection for *Ae. aegypti* strains and 6 to 9 days after infection for *Ae. albopictus* strains. For all strains, peak of mortality occurred before adult emergence, between 6 and 10 days post infection (p.i.). Figure 1 shows the percentage of cumulative mortality on day 25 after infection, corresponding to dead larvae or pupae as well as individuals that disappeared during the experiment, in the control (CTL) and infected (I) groups. In the control groups, the mortality rate was about 5% for *Ae. albopictus* strains and up to 20% for *Ae. aegypti* LHP strain at the end of the experiment. Infection with AalDV2 caused mortality in both species. However, under the same rearing and bioassay conditions, *Ae. aegypti* strains showed a higher cumulative mortality rate than *Ae. albopictus* strains (Table S1, p<0.001) suggesting that AalDV2 is more pathogenic for this species.

**Figure 1.**
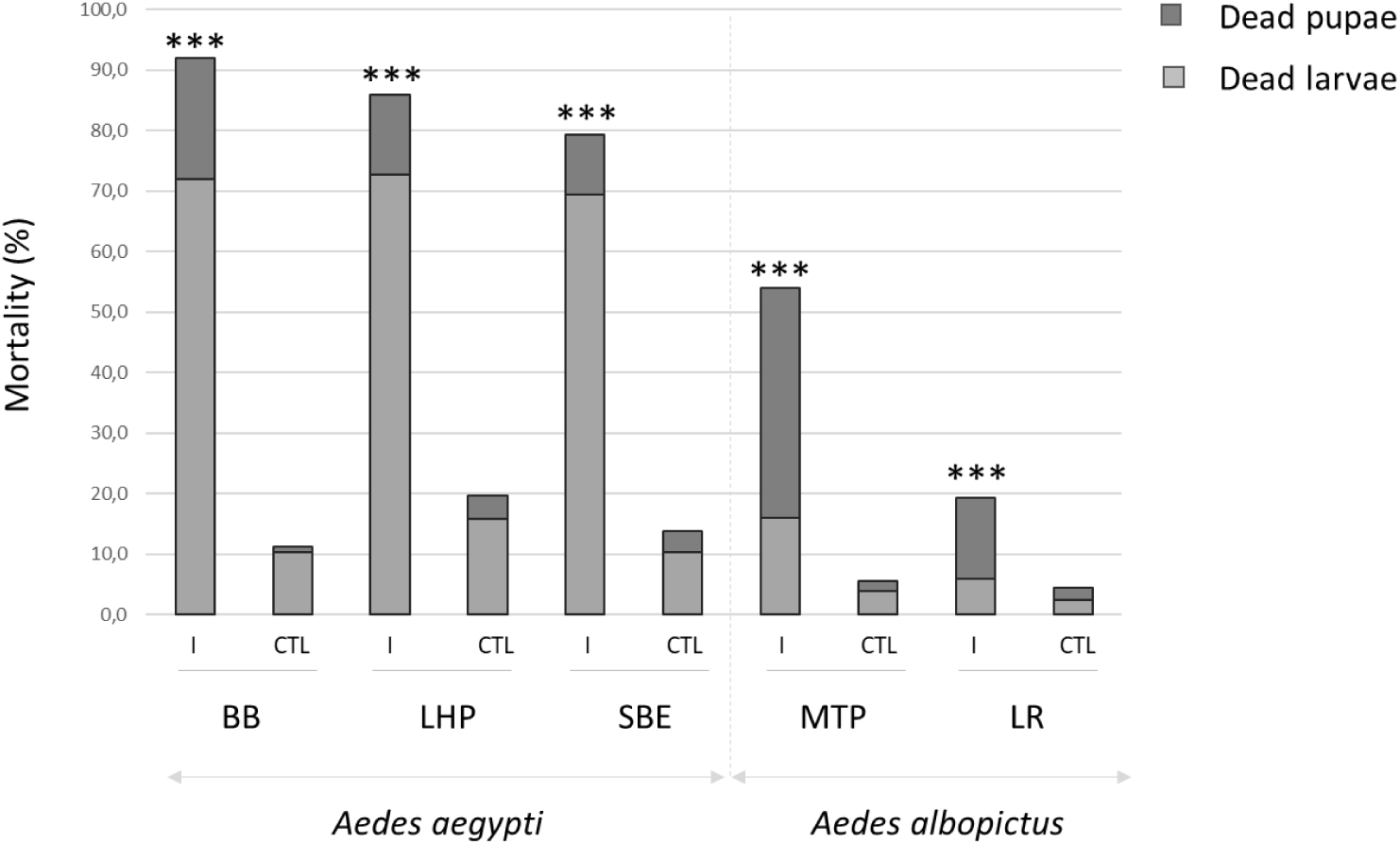
Percentage of cumulative mortality, corresponding to dead larvae or pupae, 25 days following AalDV2 infection of first instar mosquito larvae from different strains of *Ae. albopictus* (MTP, Montpellier and LR, *La Réunion*) and *Ae. aegypti* (LHP, Long-Hoà, BB, Bora-Bora and SBE, Benin). Larvae (N=3 replicates of 150 larvae per experimental condition) were infected, or not, with 3×10^11^ veg/ml of AalDV2. Binomial linear mixed effect models were used to compared the impact of AalDV2 in infected groups (I) compared to control groups (CTL). ***, p-value<0,001.

Differences in intra-specific susceptibility to the virus were also observed. In *Ae. aegypti*, the cumulative mortality of larvae and pupae infected with AalDV2 was up to 92% for the BB strain, 86% for the LHP strain and 79% for the SBE strain. Mortality rates of controls without infection were similar (Table S2, p=0.247) for BB and SBE strains, and higher than those observed for LHP (p<0.001). Viral infection increased mortality of all strains (p<0.001), but more for BB than for LHP and SBE (p<0.001). In *Ae. albopictus* strains, we observed 55% mortality in AalDV2-infected mosquitoes for the MTP strain and 19% for the LR strain. The impact of the virus on mortality was significant (Table S3, p<0.001) and greater for the MTP strain than for the LR strain (p<0.001). Mortality occurred at different stages of mosquito development. Larval mortality in *Ae. aegypti* was 73% for the LHP, 72% for the BB and 69% for the SBE strains; compared to 16% for the MTP and 6% for the LR strains of *Ae. albopictus*. Pupal mortality in *Ae. albopictus* was 38% for the MTP and 13% for the LR strains; compared to 13% for the LHP, 10% for the SBE and 20% for the BB strains of *Ae. aegypti*. Overall, there were significant differences in the pathogenicity of AalDV2 between the different strains of *Aedes* mosquitoes. Mortality was lower and occurred later for the tested strains of *Ae. albopictus* than for the *Ae. aegypti* strains.

#### Infection rate and virus concentration in dead mosquitoes

Dead mosquitoes were collected daily and analysed by qPCR for detection and quantification. In the infected groups, the virus was detected in all dead mosquitoes both in the larval and pupal stages (100%, N=1,322). Control mosquito were negative for any densoviruses infection. Moreover, no signals were detected in the control mosquito samples that died during the experiment (N=234). Figure 2 shows the viral dose quantified in dead larvae and pupae for each strain tested of *Ae. albopictus* and *Ae. aegypti*. Viral doses were slightly higher in larvae than pupae, except for the LR strain of *Ae. albopictus* where pupae were significantly more infected than larvae (Wilcoxon test, W=645, p<0,0001).

**Figure 2.**
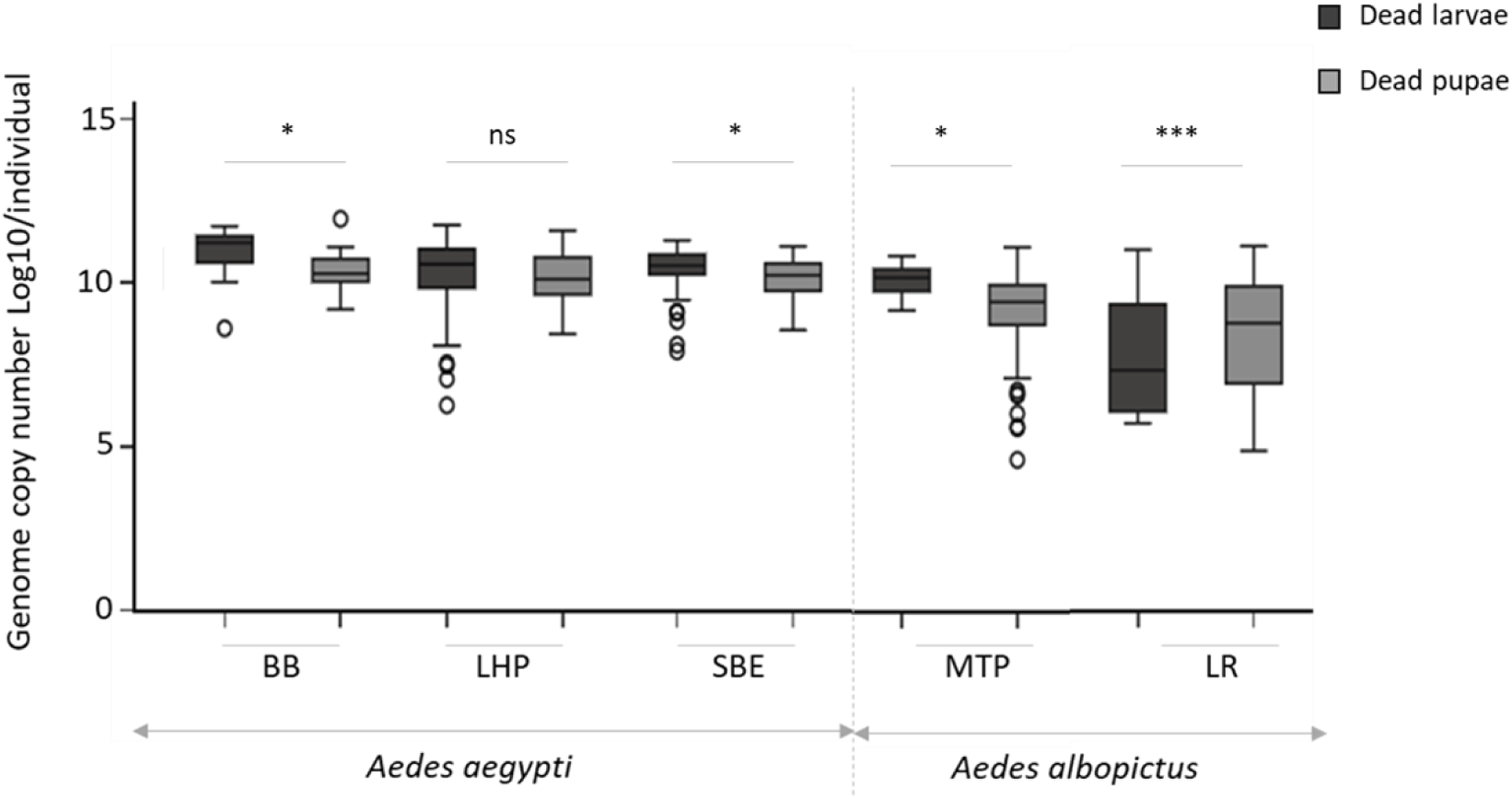
Viral dose in dead larvae or pupae of AalDV2-infected mosquitoes for each tested strain of *Ae. aegypti* (LHP, Long-Hoà, BB, Bora-Bora and SBE, Benin) and *Ae. albopictus* (MTP, Montpellier and LR, *La Réunion*). Larvae (N=3 replicates of 150 larvae per experimental condition) were infected, or not, with 3×10^11^ veg/ml of AalDV2. Viral doses of dead larvae and dead pupae are expressed as log 10 of the number of AalDV2 genomes quantified (gev) by qPCR in each individuals. Wilcoxon W test was used to compared virus titer in larvae compared to pupae for each strain. ***, p-value<0,001; *, p-value<0,05; n.s., not statistically significant.

Table 1 shows the mean viral copy number of the AalDV2 for each strain tested according to stage of development. In *Ae. aegypti* larvae, viral titers reached 5.14E+11 gev/individual in BB strain, 1.49E+11 gev/individual in LHP strain and 1.33E+11 gev/individual in SBE strains with an average of 1.73E+11 ± 6.72E+10 gev/individual for BB strain, 8.89E+10 ± 4.51E+10 gev/individual in LHP strain and 4.40E+10 ± 8.51E+09 gev/individual in SBE strain. In larvae of *Ae. albopictus*, viral titers were less important and reached 6.81E+10 gev/individual in MTP strain and 1.12E+09 gev/individual in LR strain with an average of 1.85E+10 ± 5.7E+09 gev/individual in MTP strain and 1.20E+08 ± 1.98E+08 gev/individual in LR strain. In pupae, viral titer was on average significantly lower than in larvae, although we observed very high titer in some individuals, including in *Ae. albopictus*. Indeed, the viral titers reached 8.71E+11 gev/individual in the BB strain, 3.80E+11 gev/individual in the LHP strain, 1.26E+ 11 gev/individual in the SBE strain, 1.27E+11 gev/individual in the strain MTP and 1.30E+11 gev/individual in the LR strain. On average, we obtain 7.60E+10 ± 8.76E+10 gev in the BB strain, 5.03E+10 ± 4.15E+10/individual in the LHP strain, 2.44E+10 ± 9,98E+09 gev/individual in the SBE strain of *Ae. aegypti*. In pupae of *Ae. albopictus*, viral titers were less important excepted for LR strain. We obtained an average of 9.83E+09 ± 4.6E+09 gev/individual in MTP strain and 1.35E+10 ± 9.86E+09 gev/individual in LR strain.

**Table 1.** Mean viral copy number of AalDV2 in dead larvae and pupae for each tested strain of *Ae. aegypti* (LHP, Long-Hoà, BB, Bora-Bora and SBE, Benin) and *Ae. albopictus* (MTP, Montpellier and LR, *La Réunion*). Larvae (N=3 replicates of 150 larvae per experimental condition) were infected or not, with 3×10^11^ veg/ml of AalDV2. Viral doses of dead larvae and pupae are expressed as the number of AalDV2 genomes quantified by qPCR in each individuals (gev/individual).

|  | Virus titer in dead larvae<br>(gev/individual) | Virus titer in dead pupae<br>(gev/individual) |
| --- | --- | --- |
| BB<sup>1</sup> | $1.73E+11 \pm 6.72E+10$ | $7.60E+10 \pm 8.76E+10$ |
| LHP<sup>1</sup> | $8.89E+10 \pm 4.51E+10$ | $5.03E+10 \pm 4.15E+10$ |
| SBE<sup>1</sup> | $4.40E+10 \pm 8.51E+09$ | $2.44E+10 \pm 9.98E+09$ |
| MTP<sup>2</sup> | $1.85E+10 \pm 5.87E+09$ | $9.83E+09 \pm 4.60E+09$ |
| LR<sup>2</sup> | $1.20E+08 \pm 1.98E+08$ | $1.35E+10 \pm 9.86E+09$ |

1: *Aedes aegypti*; 2: *Aedes albopictus*

### Impact of densovirus infection on mosquitoes cannibalism

At the end of the experiment, all individuals were counted. Missing individuals between the start (n=150/replicate) and the end of the trial were considered as consumed (cannibalized) by the others. Under these conditions of infection and rearing, *Ae. albopictus* strains are significantly less prone to cannibalism after DVs infection than *Ae. aegypti* strains (Table S4, p<0.001). As shown in Figure 3, in the *Ae. albopictus* strains, we observed little cannibalism with only 8.2% in the MTP strain (Table S6, p<0.05) and 1.3% in the LR strain (p=0.399). We frequently observed high cannibalism in infected groups of *Ae. aegypti*. Cannibalism of AalDV2-infected mosquitoes reached 40.7% in the BB strain, 33.6% in the LHP strain and 38.5% in the SBE strain. Viral infection increased cannibalism in all strains of *Ae. aegypti* (Table S5, p<0.001), but more so in the BB and SBE strains than in the LHP strain (p<0.001).

**Figure 3.**
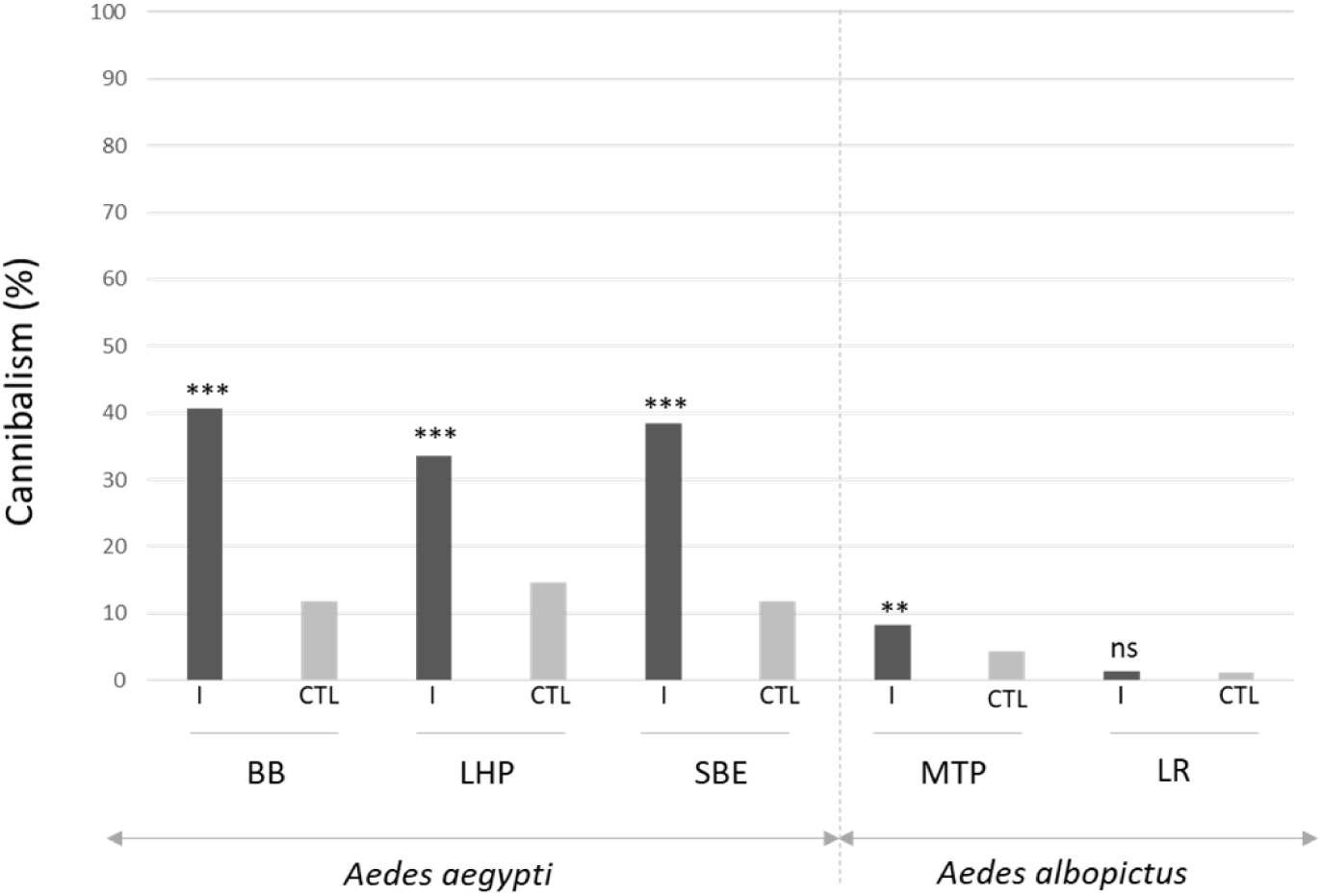
Larval cannibalism rate on day 25 after AalDV2 infection of first instar mosquito larvae for the different strains tested of *Ae. albopictus* (MTP, Montpellier and LR, *La Réunion*) and *Ae. aegypti* (LHP, Long-Hoà, BB, Bora-Bora and SBE, Benin). Larvae (N=3 replicates of 150 larvae per experimental condition) were infected or not, with 3×10^11^ veg/ml of AalDV2. Binomial linear mixed effect models were used to compared the impact of AalDV2 on cannibalism in infected groups (I) compared control groups (CTL). ***, p-value<0,001; **, p-value<0,01; n.s., not statistically significant.

### Densovirus infection on surviving adults

#### Prevalence and virus concentration

Surviving emerged adults (males and females) were collected daily and analyzed by qPCR for AalDV2 detection and quantification. The infection rate in surviving adults was very high: the virus was still present after the emergence of adults at a significant level. As shown in figure 4 the prevalence of AalDV2 in surviving adults was higher in *Ae. albopictus* strains than in *Ae. aegypti* strains, with comparable prevalence rates between MPT (90.3%) and LR (85.7%). In *Ae. aegypti*, surviving adults of the SBE strain are significantly less infected than the two other strains (Table S7, p<0.01). Thus, we observed that 62.6% (72 of 115) of surviving adults of the SBE strain, 67.6% (23 of 34) of the LHP strain, and 79.6% (43 of 54) of the BB strain of surviving adults were infected with AalDV2. The overall frequency of infection did not vary by gender. In *Ae. albopictus*, 87.9% of females and 92.7% of males for the MTP strain were infected with AalDV2 compared to 82% of females and 92.7% of males for the LR strain. In *Ae. aegypti*, the prevalence rates of AalDV2 infection were 60% of females and 78.6% of males for the LHP strain, 66.2% of females and 56.1% of males for the SBE strain, and 74.4% of females and 93.3% of males for the BB strain.

**Figure 4.**
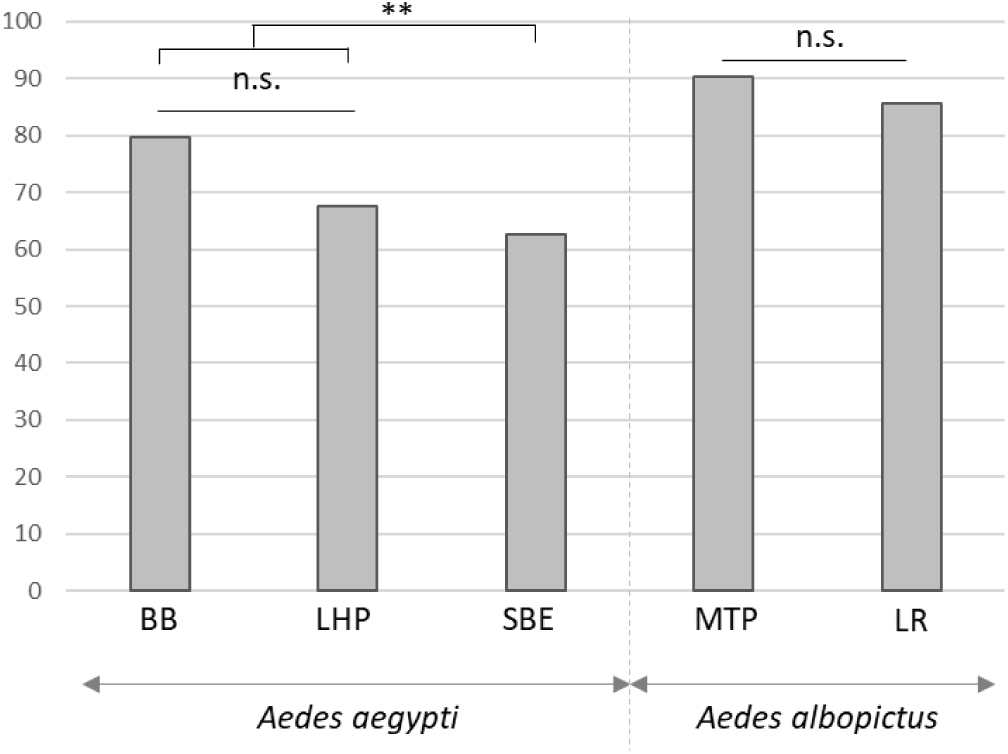
Percentage of infected adults surviving after AalDV2 infection of first instar mosquito larvae for the different strains tested of *Ae. albopictus* (MTP, Montpellier and LR, *La Réunion*) and *Ae. aegypti* (LHP, Long-Hoà, BB, Bora-Bora and SBE, Benin). Larvae (N=3 replicates of 150 larvae per experimental condition) were infected or not, with 3×10^11^ veg/ml of AalDV2. Binomial linear mixed effect models were used to compared the impact of AalDV2 in different strains. **, p-value<0,01; n.s., not statistically significant.

Figure 5 shows the viral dose quantified in surviving infected adults for each strain of *Ae. albopictus* and *Ae. aegypti*.

**Figure 5.**
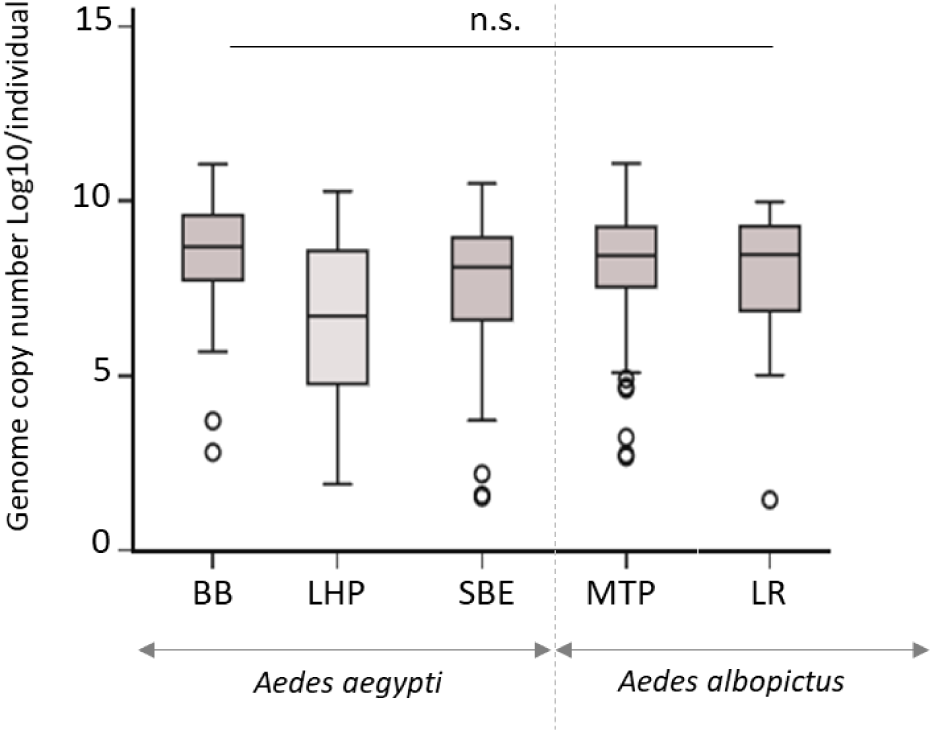
Viral dose in surviving adults of AalDV2-infected mosquitoes for each tested strain of *Ae. Albopictus* (MTP, Montpellier and LR, *La Réunion*) and *Ae. aegypti* (LHP, Long-Hoà, BB, Bora-Bora and SBE, Benin). Larvae (N=3 replicates of 150 larvae per experimental condition) were infected or not, with 3×10^11^ veg/ml of AalDV2. Viral doses of infected surviving adult are expressed as log 10 of the number of AalDV2 genomes quantified (gev) by qPCR in each individuals. Wilcoxon W test was used to compare the viral titer in adults for each strain. n.s., not statistically significant.

The viral dose in surviving infected adults was not significantly different between the different strains of *Ae. albopictus* and *Ae. aegypti* (Table 2). In *Ae. aegypti*, viral titers reached 3.61E+10 gev/individual for the BB strain, 1.84E+10 gev/individual for the LHP strain and 3.19E+10 gev/individual for the SBE strains with an average of 3.45E+09 ± 2.04E+09 gev/individual for the BB strain, 2.23E+09 ± 2.17E+09 gev/individual for the LHP strain and 2.02E+09 ± 1.30E+09 gev/individual for the SBE strain. Similarly, in *Ae. albopictus*, viral titers reached 3.42E+10 gev/individual for the MTP strain and 3.41E+10 gev/individual for the LR strain with an average of 3.12E+09 ± 1.37E+09 gev/individual for the MTP strain and 2.92E+09 ± 1.21E+09 gev/individual for the LR strain. The viral dose was not significantly different in surviving adults by gender.

**Table 2.**
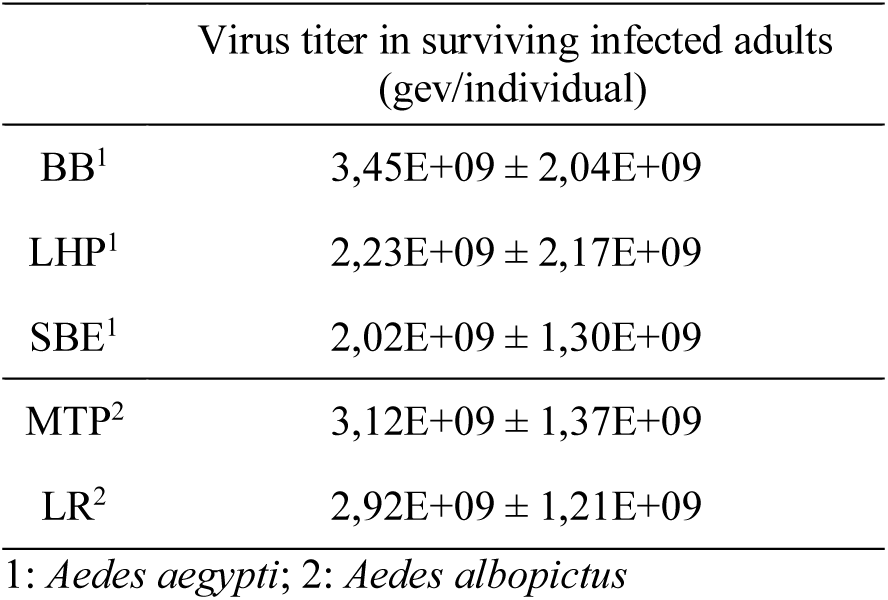
Mean viral copy number of AalDV2 in infected surviving adults for each tested strain of *Ae. aegypti* (LHP, Long-Hoà, BB, Bora-Bora and SBE, Benin) and *Ae. albopictus* (MTP, Montpellier and LR, *La Réunion*). Larvae (N=3 replicates of 150 larvae per experimental condition) were infected or not, with 3×10^11^ veg/ml of AalDV2. Viral doses of infected adult are expressed as the number of AalDV2 genomes quantified by qPCR in each individuals (gev/individual).

#### Sex ratio bias among emerging adults

The surviving emerged adults were sorted by sex. Table 3 shows the sex ratio between males and females by infection status and species. Imbalances in sex ratios were observed in the infected groups compared to control groups for SBE (p<0,05) and BB (p<0,001) strains of *Ae. aegypti* but not for LHP strain (p=0,1). No differences were observed in *Ae. albopictus* MTP (p=0,81) and LR (p=0,83) strains.

**Table 3.**
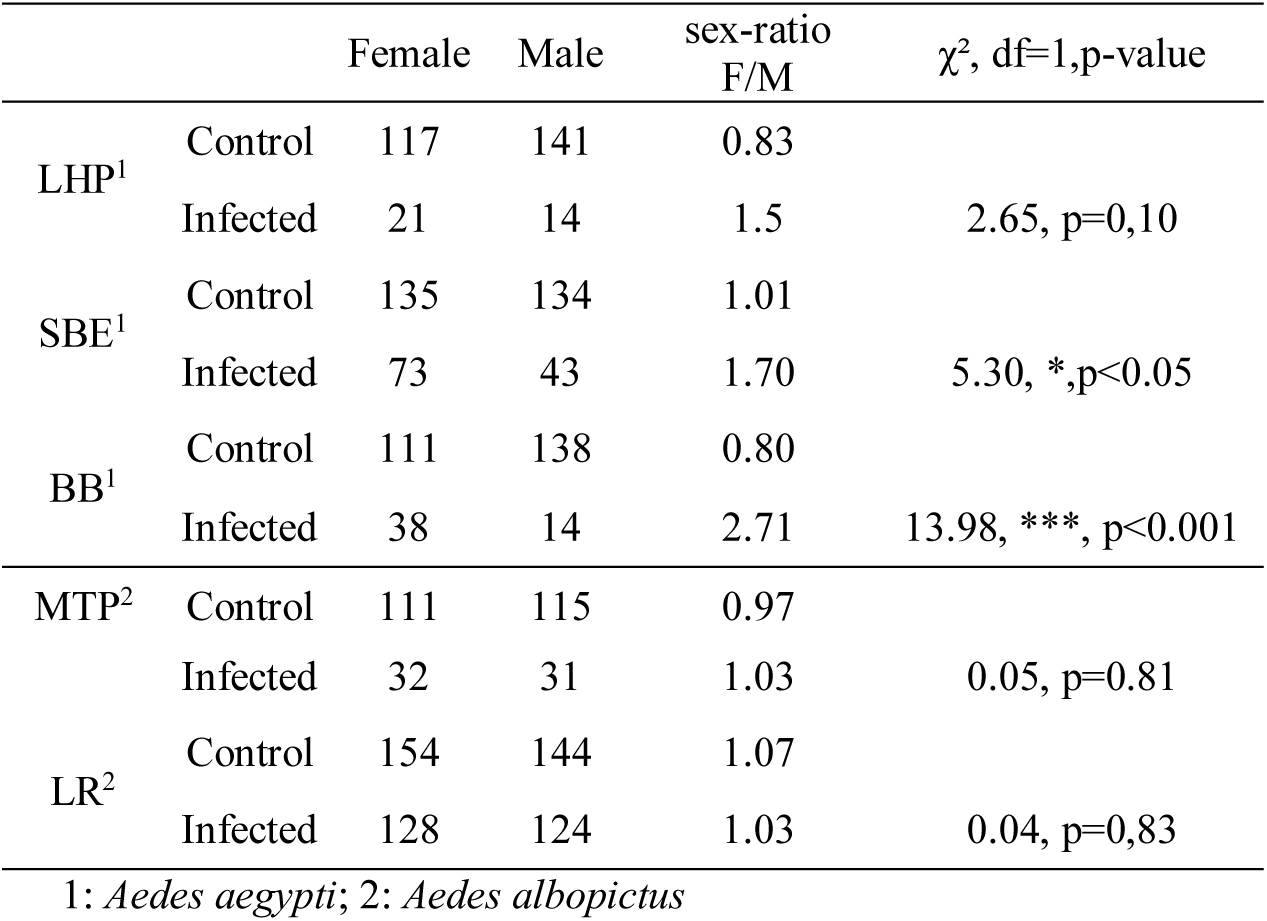
Sex ratio (female number / male number) in control and infected groups for each strains of *Ae. aegypti* (LHP, SBE, BB) and *Ae. albopictus* (MTP, LR) (*, p<0,05; ***, p<0,001; Chi-squared test).

## Discussion

Originally established from homogenates of mosquito larvae^47^, *Ae. albopictus* C6/36 cells are highly permissive to many arboviruses^48^ and are widely used for screening mosquito field collections. Aedes albopictus Densovirus 2, AalDV2, was first described in a C6/36 cell sub-line during a study on arboviruses in African mosquitoes^30^. Its origin is unknown but is probably due to contamination by samples of infected mosquito collected in the field*Our results showed that not only *Ae. aegypti* but also, for the first time, *Ae. albopictus*, were both susceptible to oral infection with AalDV2. However, we showed intra-specific and inter-specific variation in infection between the five different strains of mosquitoes tested, including insecticide resistant strains. The Long-Hoà (LHP, Vietnam) strain of *Ae. aegypti* and the *La Réunion* (LR) strain of *Ae. albopictus* are both pyrethroid-resistant strains. We have shown that these two strains are as susceptible to viral infection as the other strains. Most notably, AalDV2 appeared to be more pathogenic for *Ae. aegypti* than for *Ae. albopictus*. Mortality was mainly observed at larval stage in *Ae. aegypti* strains, compared to *Ae. albopictus* strains where less than 20 % of larvae died before pupation. Viral titers of dead individuals were higher in larvae than in pupae and in *Ae. aegypti* strains than in *Ae. albopictus* strains. This suggests that clearance of the virus could occur between each larvae moult and be released into the rearing water during the moulting process. The virus appeared to replicate less in *Ae. albopictus* strains, which could explain its lower impact on this species. AalDV2 was isolated from persistently infected cell lines derived from *Ae. albopictus* and could therefore have reduced virulence for this species as a result of a co-evolution. Studies using C6/36 densoviruses on *Ae. albopictus* are poorly documented. A single study on AalDV1 has shown that this isolate seems to be very pathogenic for the Guangdong *Ae. albopictus* strain (China)^43^. However, the mortality observed since one day post-infection is questionable and may not be linked to the virus, but to environmental factors or to rearing conditions.

We also observed a variation in susceptibility to infection at the intra-specific level. Indeed, for *Ae. aegypti*, the Bora Bora strain (BB) had a statistically higher mortality rate after exposure to AalDV2 compared to the two others strains, Long-Hoà (LHP) and Benin (SBE). This difference in intra-specific susceptibility could be a consequence of the lower genetic variability of the BB strain associated with its older establishment in the insectarium compared to the two others strains. Similarly, for *Ae. albopictus*, the Montpellier strain (MTP) had a much higher mortality than the *La Réunion* strain (LR). The dynamics of infectious diseases can be affected by genetic diversity within host populations^50^ as well as by the time of colonization of laboratory populations of *Aedes* mosquitoes^51^. Previous studies using MDVs have shown a high level of mortality for the same species of mosquito exposed to different isolates of DVs, with some isolates being highly pathogenic, others more benign. For example, as they are different viral strains, AalDV1 infection resulted in a mortality rate of over 90% in 1st instar *A. aegypti* larvae^30^, while AaeDV infection results in a mortality of 75.1% for the same species^52^. Thus, after infection with AthDV, first-instar larvae of the Thai strain of *Ae. albopictus*, had a mortality of 82% compared to 51% for *Ae. aegypti*^41^. Different DVs strains such as AaeDV1, AthDV, AalDV2 and AalDV3 induced completely different levels of mortality when infecting Rexville D, Chachoengsao, and Bangkok strains of *Ae. aegypti* after 48h exposure to DVs at 2×10^10^ gev/larvae^49^. Recently, analysis of sublethal effects also showed that AalDV7 infection of *Ae. aegypti* and *Ae. albopictus* first instar larvae significantly decreased pupation and emergence rates^36^. However, the diversity of experimental designs used does not allow easy comparison of the results across studies, due to differences in infection methods, environmental conditions, viral titers, and stage of larvae infected. Some intrinsic factors such as their genetic background, or their microbiota, could influence the susceptibility of mosquitoes to DVs infection but these mechanisms need to be explored. For example, in *Bombyx mori* lepidoptera, resistance to Bombyx mori densovirus type 1 or 2 (BmDV1 & BmDV2) is controlled by recessive non-susceptibility genes, nsd-1 and -2^53,54^, which affect distinct stages of the viral infection pathway^55^. In addition, a recent study has shown that *Wolbachia pipientis* infection promotes the replication of the Aedes albopictus densovirus 1 (AalDV1) in *Aedes* cell lines in a density dependent manner^56^. Further studies are needed to determine how these laboratory results may result in increased susceptibility to DVs in natural populations of *Wolbachia*-infected *Ae. albopictus* or artificially *Wolbachia*-infected *Ae aegypti*. Furthermore, the pathogenicity, prevalence and infection rate of MDVs may also vary with on environmental factors such as temperature, or other conditions such as larval density, method of infection and duration of exposure to MDVs^45,57^. The environmental factors that influence the efficacy of MDVs have not been thoroughly studied and further research is needed before these MDVs can be eligible for operational use in mosquito control programs.

In mosquitoes, cannibalism between larval instars of the same species has been frequently observed, especially in the later instars, with food deficiency or excessively high larval densities applied under rearing conditions. Cannibalism is also an effective route of transmission for some pathogens, including DVs, when healthy larvae consume moribund infected ones. Sick larvae become lethargic in the later stages of infection and are unable to defend themselves against aggressive conspecifics. Under our laboratory conditions, *Ae. albopictus* and *Ae. aegypti* had a larval development cycle of 9-10 days at 26°C from egg-laying to emergence. During our experiment, infected larvae of *Ae. aegypti* showed a delay in development compared to unexposed larvae and especially compared to infected larvae of *Ae. albopictus*. After infection in the first-instar stage, larval development of infected *Ae. aegypti* larvae was heterogeneous compared to healthy larvae which have a more synchronized life cycle. Most cadavers collected were third and fourth instar larvae whereas first or second instar larvae were rarely found. The high incidence of larval cannibalism observed in *Ae. aegypti* mosquitoes, a species more susceptible to AalDV2, suggests that this is an important pathway for the transmission and pathogenicity of this MDVs.

Adult mortality has not been evaluated in this work, but many studies have shown that DVs infection of mosquito larvae affects the life traits of infected adults. Thus, the effects of sublethal infection of *Ae. aegypti* larvae by different isolates of DVs included extended larval development times, reduced pupal and adults weight, decreased fertility of females, and decreased adult lifespan. Although the ability to modify adult life characteristics, in particular fertility, differs between MDVs strains and mosquito species, a reduction in reproductive success could potentially lead to a decrease in mosquito density and vector capacity.

Females infected in the larval stage can transmit DVs vertically by laying infected eggs in new oviposition sites, resulting in the spread of MDVs in the mosquito population and an increased coverage and efficacy^60^. A semi-field trial have shown that adult female *Ae. aegypti* oviposition behavior led to successful AaeDV dispersal from treated breeding sites to new breeding sites in large-scale cages. However, the AaeDV titers achieved in the contaminated sites were not sufficient to reduce larval densities^40^. Further research is needed to assess for other MDVs whether this vertical transmission translates into operational efficiency in the field, which would be possible after one or more amplification cycles, as suggested by Carlson^38^. Analysis of infection rates and titers in live adults revealed that virus replication occurred in all strains for both *Aedes* species tested. Most surviving adults were positive to AaLDV2 detection after infection of first instar larvae. Infection rates in surviving adult of *Ae. albopictus* was higher than that of *Ae. aegypti* reaching 90%. Virus replication occurred in all strains for both mosquito species tested with an average viral titre of 6.5 to 11 log/ after potential exposure of first larval stage to 10 log of virus. Among the few emerging adults, the viral titer of AalDV2 infected females and males did not vary significantly. However, in some mosquito’s strains, we observed a distortion of the sex-ratio in favour of females. Butchasky et al. also observed a mortality rate in adult males infected at the larval stage with DVs three times higher than that females^61^. This selection could be of interest for spreading and maintaining the virus by vertical transmission from female to the offspring.

The results of this laboratory study provide baseline data on the susceptibility of *Ae. albopictus* and *Ae. aegypti* to AalDV2 infection. We have shown that the different strains of mosquitoes were susceptible to AalDV2 infection including for insecticides resistance strains. These results confirm even more the advantage of the isolate AalDV2 and of the DVs in general as a biological control agent. The differences in the pathogenicity of AalDV2 among *Aedes* mosquito strains draw attention to the risk associated with the development of single viral strain for use as biocontrol agents. Intra-specific variations suggest that host resistance to infection may evolve. Our work also suggest that DVs strains isolated from heterologous mosquito species may be more efficient against a given species, probably because they are less well adapted. In addition, highly pathogenic DVs that kill larvae before they reach adulthood may exert high selective pressure, which would increase the risk of resistance and decrease their efficacy over time. Thus, AalDV2 could be a potential mosquito control agent, but further research and development work is still needed, including studies on the different routes of transmission and its persistence in the environment through semi-field evaluations. The continued discovery and isolation of new MDVs will enrich the pool entomopathogenic mosquito viruses and provide a variety of choices for one or more combinations of MDVs to optimally target *Aedes* mosquitoes. The ability to produce the virus on a large-scale at low cost and in sufficient quantities is also required before innovative formulations can be developed for operational use with other methods as part of integrated vector management. The development of innovative formulations suitable for i) direct use in breeding site or ii) dissemination by the insects themselves according to the entomovectoring principle developed by the boosted SIT approach, is underway^44^.

## Methods

### Virus and cells line

We used two *Ae. albopictus* derived C6/36 cell lines^62^. The first, a chronically infected sub-line, was used as the source of the Aedes albopictus densovirus 2 (AalDV2)^30^. A second, free of any DVs, was used as a control. Both sub-lines were grown at 28°C in RPMI medium (Dutcher, France), supplemented with 10% heat-inactived fetal calf serum (Gibco™, USA), 1% non-essential amino-acids (Gibco™, USA) and 1% penicillin-streptomycin (Gibco™, USA). Cells were seeded at 7-days intervals in 25 cm^2^ flasks at 1:5 dilution. Infected cells were scraped into supernatant and kept at −20°C as virus stock. The viral concentration was estimated as described below. The titer, expressed as genome equivalent virus (gev), remained stable at 3×10^11^ gev/ml.

### Mosquitoes strains

Three different colonized strains of *Ae. aegypti* and two of *Ae. albopictus* have been used to study the pathogenicity of AalDV2 (Table 4).

**Table 4.** *Aedes albopictus* and *Ae. aegypti* mosquitoes strains. ^1^Susceptible to all insecticides class, ^2^Resistant to Pyrethroids, the main insecticide class used for Aedes vector control. * The characterization of molecular and metabolic mechanisms of insecticide resistance is underway.

| Species | Strains | Origin | Lab colonization | Status |
| --- | --- | --- | --- | --- |
| *Ae. aegypti* | BB | French Polynesia | 1980 | Susceptible<sup>1</sup> |
|  | SBE | Benin | 2008 | Susceptible<sup>1</sup> |
|  | LHP | Vietnam | 1997 | Resistant<sup>2</sup> (*kdr*) |
| *Ae. albopictus* | MTP | France | 2016 | Susceptible<sup>1</sup> |
|  | LR | Overseas France | 2016 | Resistant<sup>2*</sup> |

The reference strain of *Ae. aegypti* BB (Bora-Bora, French polynesia) and SBE (Benin) have been colonized for many years and were devoid of any phenotypic resistance to World Health Organization (WHO) susceptibility tests at diagnostic doses for the most common chemical insecticides or any known mechanism of insecticide resistance. The LHP (Long-Hoà, Vietnam), pyrethroid-resistant strain of *Ae. aegypti*, is homozygous for the knockdown resistance (*kdr*) gene^63^. In addition, we used two recently colonized *Ae. albopictus* strains from France. The *La Réunion* strain (LR), is a pyrethroid-resistant strain colonized since 2010 but whose resistance mechanism is currently under investigation. The Montpellier strain (MTP) susceptible to all insecticides has been colonized since 2016. Adult colonies are maintained in the Vectopole insectarium of IRD in Montpellier, France at 28°C, 70% humidity with a 14 h/10 h light/dark cycle and fed on pathogen free rabbit blood using a parafilm-based method and 10 % sugar solution. The larvae were reared at 28°C in 2 liters jars and fed with alevin powder. At 28°C, pupae were obtained 6-7 days after immersion of the eggs in water and imago 2 days later.

### Virus infection of mosquito larvae from *Ae. aegypti* and *Ae. albopictus*

Newly hatched first instar *Aedes* larvae were infected as follow. Mosquito eggs were allowed to hatch in tap water with a 7.5% solution of 50:50 alevin powder and rabbit pellets. Twenty-four hours after hatching, pools of 150 larvae were exposed to 3×10^11^ gev/ml of cells infected with AalDV2, theoretically corresponding to 10^10^ gev per larvae, in a total volume of 5 ml and kept without food for 48 hours. The control groups, were exposed to healthy C6/36 cells under identical conditions to those of the treatment groups. Two days after infection, larvae were transferred to 300 ml water bowls with food and observed daily until pupation. Pupae were transferred to new small cups of clear water and allowed to emerge into mosquito cages. Dead larvae or pupae, were collected daily and stored at −20°C for further investigations. The emerged adults were collected daily and sorted by sex. The larval, pupal and cumulative mortality was evaluated at the end of bioassays at day 25 after infection. The adult mortality has not been assessed. Three biological replicates of each strain were performed.

Cannibalism, defined here as the consumption of an individual of the same species, was evaluated as the number of individuals that have disappeared during the experiment. The cumulative mortality observed at the end of the experiment takes into account the number of dead larvae and pupae plus the rate of cannibalism.

### Virus detection and quantification in mosquitoes

Quantification of the virus by qPCR was performed using the LightCycler 480 System (Roche, France) and specific primers designed in the non-structural gene NS1 (q*Aal*DV2-F: 5’-TggCCAACAATTACgAACAA-3’and q*Aal*DV2-R: 5’-CTCTggAgCCgCTgTgTAAT-3’). A standard curve (10^9^ to 10^3^ viral genome copies per reaction) was generated using 10-fold serial dilutions of p*Aal*DV2, a plasmid encompassing the entire *Aal*DV2 sequence^64^. The reactions were carried out in a 10 μL reaction mixture containing 5 µl of 1X SyberGreen master mix I, 1.7 µl of DNase/RNase-free water, 0.4 µM of each primers and 2.5 µl of sample. Each sample was processed in triplicate under the following conditions: 95°C for 3’, 45 cycles of 94°C for 10 s, 60°C for 10 s and 72°C for 10 s. The data were analyzed using Light Cycler 480 software (Roche, France). Virus concentration in mosquitoes was determined individually and titers were expressed as genome equivalent virus (gev) per individual. Each individual (larvae, pupae or adult) was crushed in 50 µl of 0.1X Tris-EDTA buffer supplemented with 0.05 µM salmon sperm DNA and the suspension was clarified for 5 min at 5,000 g before quantification.

### Statistical analysis

Binomial linear mixed effect models were used to analyse the impact of AalDV2 infection on survival at intra- and inter-specific level, cannibalism and infection rates in surviving adults (response variables). The mosquito species and strain as well as the infection status were used as fixed effects and the repetitions as random effects. Fixed-effects coefficients of all models and their corresponding p-values are reported in Tables S1 to S7. Data from viral concentration in dead or surviving individuals were analysed with Wilcoxon W test. Datas from sex-ratio bias were analysed with chi-square test. By convention, results were considered statistically significant when p < .05.

### Data Availability

The datasets generated during and/or analysed during the current study are available from the corresponding author on reasonable request.

## Acknowledgments

This study was funded by EU ERC CoG - 682387 REVOLINC. The contents of this publication are the sole responsibility of the authors and do not necessarily reflect the views of the European Commission. We thank the Institute of Research and Development (IRD) and the Vectopole Sud network (http://www.vectopole-sud.fr/) for providing the infrastructure needed for insect experimentation. Thanks to the qPHD Platform (Montpellier GenomiX).

## Author Contributions

A.P. designed and performed the experiments, analysed the data. T.B., J.B., M.O., A.G.G, C.L., and F.C. assisted in data analysis. M.R., C.G., and B.S, provided technical support to experiments. A.P., J.B., T.B. and F.C. wrote the first draft of the manuscript. All authors reviewed the manuscript.

## References

1. Weaver, S. C., Charlier, C., Vasilakis, N. & Lecuit, M. Zika, Chikungunya, and Other Emerging Vector-Borne Viral Diseases. Annu. Rev. Med. 69, 395–408 (2018).

2. Kilpatrick, A. M. & Randolph, S. E. Drivers, dynamics, and control of emerging vector-borne zoonotic diseases. Lancet 380, 1946–1955 (2012).

3. Koureas, M., Tsakalof, A., Tsatsakis, A. & Hadjichristodoulou, C. Systematic review of biomonitoring studies to determine the association between exposure to organophosphorus and pyrethroid insecticides and human health outcomes. Toxicol. Lett. 210, 155–168 (2012).

4. Peterson Robert K.D., Macedo Paula A. & Davis Ryan S. A Human-Health Risk Assessment for West Nile Virus and Insecticides Used in Mosquito Management. Environ. Health Perspect. 114, 366–372 (2006).

5. Han, W., Tian, Y. & Shen, X. Human exposure to neonicotinoid insecticides and the evaluation of their potential toxicity: An overview. Chemosphere 192, 59–65 (2018).

6. Hernández, A. F. et al. Toxic effects of pesticide mixtures at a molecular level: Their relevance to human health. Toxicology 307, 136–145 (2013).

7. Sanchez-Bayo, F. P. Insecticides Mode of Action in Relation to Their Toxicity to Non-Target Organisms. J. Environ. Anal. Toxicol. s4, (2012).

8. Rivero, A., Vézilier, J., Weill, M., Read, A. F. & Gandon, S. Insecticide Control of Vector-Borne Diseases: When Is Insecticide Resistance a Problem? PLOS Pathog. 6, e1001000 (2010).

9. Hemingway, J., Hawkes, N. J., McCarroll, L. & Ranson, H. The molecular basis of insecticide resistance in mosquitoes. Insect Biochem. Mol. Biol. 34, 653–665 (2004).

10. Liu, N., Xu, Q., Zhu, F. & Zhang, L. Pyrethroid resistance in mosquitoes. Insect Sci. 13, 159–166 (2006).

11. Rivero, A., Vézilier, J., Weill, M., Read, A. F. & Gandon, S. Insecticide Control of Vector-Borne Diseases: When Is Insecticide Resistance a Problem? PLOS Pathog. 6, e1001000 (2010).

12. Dusfour, I. et al. Management of insecticide resistance in the major Aedes vectors of arboviruses: Advances and challenges. PLoS Negl. Trop. Dis. 13, e0007615 (2019).

13. Faraji, A. & Unlu, I. The Eye of the Tiger, the Thrill of the Fight: Effective Larval and Adult Control Measures Against the Asian Tiger Mosquito, Aedes albopictus (Diptera: Culicidae), in North America. J. Med. Entomol. 53, 1029–1047 (2016).

14. Chan, K. L., Ho, B. C. & Chan, Y. C. Aedes aegypti (L.) and Aedes albopictus (Skuse) in Singapore City. Bull. World Health Organ. 44, 629–633 (1971).

15. Sansinenea, E. Bacillus thuringiensis biotechnology. (Springer, 2012).

16. Mulla, M. S., Darwazeh, H. A. & Zgomba, M. Effect of some environmental factors on the efficacy of Bacillus sphaericus 2362 and Bacillus thuringiensis (H-14) against mosquitoes. Bull. Soc. Vector Ecol. 15, 166–175 (1990).

17. Marina, C. F., Arredondo-Jiménez, J. I., Castillo, A. & Williams, T. Sublethal effects of iridovirus disease in a mosquito. Oecologia 119, 383–388 (1999).

18. Delhon, G. et al. Genome of Invertebrate Iridescent Virus Type 3 (Mosquito Iridescent Virus). J. Virol. 80, 8439–8449 (2006).

19. Linley, J. R. & Nielsen, H. T. Transmission of a mosquito iridescent virus in Aedes taeniorhynchus: Laboratory experiments. J. Invertebr. Pathol. 12, 7–16 (1968).

20. Carlson, J., Suchman, E. & Buchatsky, L. Densoviruses for Control and Genetic Manipulation of Mosquitoes. in Advances in Virus Research vol. 68 361–392 (Academic Press, 2006).

21. Johnson, R. M. & Rasgon, J. L. Densonucleosis viruses (‘densoviruses’) for mosquito and pathogen control. Curr. Opin. Insect Sci. 28, 90–97 (2018).

22. Grenet, A.-S. G. et al. Les densovirus : une «massive attaque» chez les arthropodes. Virologie 19, 19–31 (2015).

23. Hewson, I. et al. Densovirus associated with sea-star wasting disease and mass mortality. Proc. Natl. Acad. Sci. 111, 17278–17283 (2014).

24. Cotmore, S. F. et al. ICTV Virus Taxonomy Profile: Parvoviridae. J. Gen. Virol. 100, 367–368 (2019).

25. Afanasiev, B. N., Galyov, E. E., Buchatsky, L. P. & Kozlov, Y. V. Nucleotide sequence and genornic organization of aedes densonucleosis virus. Virology 185, 323–336 (1991).

26. Sivaram, A. et al. Isolation and Characterization of Densonucleosis Virus from *Aedes aegypti* Mosquitoes and Its Distribution in India. Intervirology 52, 1–7 (2009).

27. Chen, S. et al. Genetic, biochemical, and structural characterization of a new densovirus isolated from a chronically infected Aedes albopictus C6/36 cell line. Virology 318, 123–133 (2004).

28. Zhai, Y. -g. et al. Isolation and characterization of the full coding sequence of a novel densovirus from the mosquito Culex pipiens pallens. J. Gen. Virol. 89, 195–199 (2008).

29. Ren, X., Hoiczyk, E. & Rasgon, J. L. Viral Paratransgenesis in the Malaria Vector Anopheles gambiae. PLoS Pathog. 4, e1000135 (2008).

30. Jousset, F.-X., Barreau, C., Boublik, Y. & Cornet, M. A Parvo-like virus persistently infecting a C6/36 clone of Aedes albopictus mosquito cell line and pathogenic for Aedes aegypti larvae. Virus Res. 29, 99–114 (1993).

31. Afanasiev, B. N. & Carlson, J. O. A new mosquito densovirus from Peru: genomic sequence and in vitro growth characteristics of wild type and hybrid viruses. (2003).

32. O’Neill, S. L. et al. Insect densoviruses may be widespread in mosquito cell lines. J. Gen. Virol. 76, 2067–2074 (1995).

33. Jousset, F.-X., Baquerizo, E. & Bergoin, M. A new densovirus isolated from the mosquito Culex pipiens (Diptera: Culicidae). Virus Res. 67, 11–16 (2000).

34. Sangdee, K. & Pattanakitsakul, S. New genetic variation of Aedes albopictus Densovirus isolated from mosquito C6/36 cell line. Southeast Asian J Trop Med Public Health 43, 12 (2012).

35. O’Neill, S. L. et al. Insect densoviruses may be widespread in mosquito cell lines. J. Gen. Virol. 76, 2067–2074 (1995).

36. Li, J. et al. A Novel Densovirus Isolated From the Asian Tiger Mosquito Displays Varied Pathogenicity Depending on Its Host Species. Front. Microbiol. 10, (2019).

37. Kittayapong, P., Baisley, K. J. & O’Neill, S. L. A mosquito densovirus infecting Aedes aegypti and Aedes albopictus from Thailand. Am. J. Trop. Med. Hyg. 61, 612–617 (1999).

38. Carlson, J., Suchman, E. & Buchatsky, L. Densoviruses for Control and Genetic Manipulation of Mosquitoes. in Advances in Virus Research vol. 68 361–392 (Academic Press, 2006).

39. Barreau, C., Jousset, F. X. & Bergoin, M. Venereal and vertical transmission of the Aedes albopictus parvovirus in Aedes aegypti mosquitoes. Am. J. Trop. Med. Hyg. 57, 126–131 (1997).

40. De Valdez, M. R. W., Suchman, E. L., Carlson, J. O. & Black, W. C. A Large Scale Laboratory Cage Trial of Aedes Densonucleosis Virus (AeDNV). J. Med. Entomol. 47, 392–399 (2010).

41. Kittayapong, P., Baisley, K. J. & O’Neill, S. L. A mosquito densovirus infecting Aedes aegypti and Aedes albopictus from Thailand. Am.J.Trop.Med.Hyg. 61, 612–617.

42. Altinli, M. et al. Sharing cells with Wolbachia: the transovarian vertical transmission of Culex pipiens densovirus. Environ. Microbiol. 21, 3284–3298 (2019).

43. Wei, W. et al. The pathogenicity of mosquito densovirus (C6/36DNV) and its interaction with dengue virus type II in Aedes albopictus. Am. J. Trop. Med. Hyg. 75, 1118–1126 (2006).

44. Bouyer, J., Chandre, F., Gilles, J. & Baldet, T. Alternative vector control methods to manage the Zika virus outbreak: more haste, less speed. Lancet Glob. Health 4, e364 (2016).

45. Barreau, C., Jousset, F.-X. & Bergoin, M. Pathogenicity of theAedes albopictusParvovirus (AaPV), a Denso-like Virus, forAedes aegyptiMosquitoes. J. Invertebr. Pathol. 68, 299–309 (1996).

46. Barreau, C., Jousset, F.-X. & Cornet, M. An efficient and easy method of infection of mosquito larvae from virus-contaminated cell cultures. J. Virol. Methods 49, 153–156 (1994).

47. Igarashi, A. Isolation of a Singh’s Aedes albopictus Cell Clone Sensitive to Dengue and Chikungunya Viruses. J. Gen. Virol. 40, 531–544 (1978).

48. Brackney, D. E. et al. C6/36 Aedes albopictus Cells Have a Dysfunctional Antiviral RNA Interference Response. PLoS Negl. Trop. Dis. 4, e856 (2010).

49. Hirunkanokpun, S., Carlson, J. O. & Kittayapong, P. Evaluation of Mosquito Densoviruses for Controlling Aedes aegypti (Diptera: Culicidae): Variation in Efficiency due to Virus Strain and Geographic Origin of Mosquitoes. Am. J. Trop. Med. Hyg. 78, 784–790 (2008).

50. Ostfeld, R. S. & Keesing, F. Effects of Host Diversity on Infectious Disease. Annu. Rev. Ecol. Evol. Syst. 43, 157–182 (2012).

51. Lambrechts, L., Scott, T. W. & Gubler, D. J. Consequences of the Expanding Global Distribution of Aedes albopictus for Dengue Virus Transmission. PLoS Negl. Trop. Dis. 4, e646 (2010).

52. 1. 52. Ledermann, J. P., Suchman, E. L., Black, W. C. & Carlson, J. O. Infection and Pathogenicity of the Mosquito Densoviruses AeDNV, HeDNV, and APeDNV in Aedes aegypti Mosquitoes (Diptera: Culicidae). J. Econ. Entomol. 97, 1828–1835 (2004).

53. Ogoyi, D. O. et al. Linkage and mapping analysis of a non-susceptibility gene to densovirus (nsd-2) in the silkworm, Bombyx mori. Insect Mol. Biol. 12, 117–124 (2003).

54. Watanabe, H. & Maeda, S. Genetically determined nonsusceptibility of the silkworm, Bombyx mori, to infection with a densonucleosis virus (Densovirus). J. Invertebr. Pathol. 38, 370–373 (1981).

55. Rudolf, V. H. W. & Antonovics, J. Disease transmission by cannibalism: rare event or common occurrence? Proc. R. Soc. B Biol. Sci. 274, 1205–1210 (2007).

56. Parry, R., Bishop, C., De Hayr, L. & Asgari, S. Density-dependent enhanced replication of a densovirus in Wolbachia-infected Aedes cells is associated with production of piRNAs and higher virus-derived siRNAs. Virology 528, 89–100 (2019).

57. Rwegoshora, R. T., Baisley, K. J. & Kittayapong, P. Seasonal and spatial variation in natural densovirus infection in Anopheles minimus s.l. in Thailand. Southeast Asian J Trop Med Public Health 31, 7 (2000).

58. Edgerly, J. S., Willey, M. S. & Livdahl, T. Intraguild Predation Among Larval Treehole Mosquitoes, Aedes albopictus, Ae. aegypti, and Ae. triseriatus (Diptera: Culicidae), in Laboratory Microcosms. J. Med. Entomol. 36, 394–399 (1999).

59. Weissman, D. B., Gray, D. A., Pham, H. T. & Tijssen, P. Billions and billions sold: Pet-feeder crickets (Orthoptera: Gryllidae), commercial cricket farms, an epizootic densovirus, and government regulations make for a potential disaster. Zootaxa 3504, 67 (2012).

60. Ren, X. & Rasgon, J. L. Potential for the Anopheles gambiae Densonucleosis Virus To Act as an “Evolution-Proof” Biopesticide. J. Virol. 84, 7726–7729 (2010).

61. Buchatsky, L. P. Densonucleosis of blood sucking mosquitoes. Auq Org 6, 14–150.

62. Igarashi, A. Isolation of a Singh’s Aedes albopictus Cell Clone Sensitive to Dengue and Chikungunya Viruses. J. Gen. Virol. 40, 531–544 (1978).

63. Brengues, C. et al. Pyrethroid and DDT cross-resistance in Aedes aegypti is correlated with novel mutations in the voltage-gated sodium channel gene. Med. Vet. Entomol. 17, 87–94 (2003).

64. Boublik, Y., Jousset, F.-X. & Bergoin, M. Complete Nucleotide Sequence and Genomic Organization of the Aedes albopictus Parvovirus (AaPV) Pathogenic for Aedes aegypti Larvae. Virology 200, 752–763 (1994).

