## Supplementary tables for "Mosquito densoviruses: the revival of a biological control agent against urban *Aedes* vectors of arboviruses"

**Table S1** - Fixed-effects coefficients of a mixed-effect binomial model of the densovirus infection effect on the *Aedes* species survival. *Aedes aegypti* uninfected batches are considered as the reference level in the model (30 observations, 3 repeats).

| Fixed effects | Value | Std. Error | z-value | p-value |
|---|---|---|---|---|
| Intercept | 1.7554 | 0.1284 | 13.671 | <2e-16 |
| *Ae. albopictus* | 1.1753 | 0.1752 | 6.707 | 1.99 e-11 |
| Infection | -3.5597 | 0.1141 | -31.199 | <2e-16 |
| *Ae. albopictus* : infection | 1.1211 | 0.2082 | 5.384 | 7.29e-08 |

**Table S2** - Fixed-effects coefficients of a mixed-effect binomial model of the densovirus infection effect on the *Ae. aegypti* strains survival. Uninfected batches of Bora Bora strain (BB) are considered as the reference level in the model (18 observations, 3 repeats).

| Fixed effects | Value | Std. Error | z-value | p-value |
|---|---|---|---|---|
| Intercept | 2.0723 | 0.1740 | 11.909 | <2e-16 |
| LHP strain | -0.6603 | 0.1950 | -3.386 | 0.000708 |
| SBE strain | -0.2369 | 0.2049 | -1.156 | 0.247713 |
| Infection | -4.4289 | 0.2279 | -19.437 | <2e-16 |
| LHP strain : infection | 1.2006 | 0.2958 | 4.059 | 4.92e-05 |
| SBE strain*:* infection | 1.2458 | 0.2946 | 4.229 | 2.35e-05 |

**Table S3** - Fixed-effects coefficients of a mixed-effect binomial model of the densovirus infection effect on the *Ae. albopictus* strains survival. Uninfected batches of *La Réunion* strain (LR) are considered as the reference level in the model (12 observations, 3 repeats).

| Fixed effects | Value | Std. Error | z-value | p-value |
|---|---|---|---|---|
| Intercept | 3.0801 | 0.2938 | 10.485 | <2e-16 |
| MTP strain | -0.2325 | 0.3156 | -0.737 | 0.461342 |
| Infection | -1.6726 | 0.2715 | -6.160 | 7.27e-10 |
| MTP strain : infection | -1.3075 | 0.3590 | -3.642 | 0.000271 |

**Table S4** - Fixed-effects coefficients of a mixed-effect binomial model of densovirus infection effect on the *Aedes* species cannibalism. *Aedes aegypti* uninfected batches are considered as the reference level in the model (30 observations, 3 repeats).

| Fixed effects | Value | Std. Error | z-value | p-value |
|---|---|---|---|---|
| Intercept | -2.2266 | 0.1896 | -11.743 | <2e-16 |
| *Ae. albopictus* | -1.4444 | 0.2355 | -6.132 | 8.66 e-10 |
| Infection | 1.8322 | 0.1106 | 16.575 | <2e-16 |
| *Ae. albopictus* : infection | -1.2209 | 0.2985 | -4.090 | 4.31e-05 |

**Table S5** - Fixed-effects coefficients of a mixed-effect binomial model of the densovirus infection effect on the *Ae. aegypti* strains cannibalism. Uninfected batches of LHP strain are considered as the reference level in the model (18 observations, 3 repeats).

| Fixed effects | Value | Std. Error | z-value | p-value |
|---|---|---|---|---|
| Intercept | -1.8258 | 0.1922 | -9.498 | <2e-16 |
| BB strain | -0.6731 | 0.2283 | -2.949 | 0.00319 |
| SBE strain | -0.5828 | 0.2249 | -2.592 | 0.00955 |
| Infection | 0.9409 | 0.1795 | 5.242 | 1.59e-07 |
| BB strain : infection | 1.4526 | 0.2713 | 5.354 | 8.58e-08 |
| SBE strain: infection | 1.2400 | 0.2706 | 4.582 | 4.60e-06 |

**Table S6** - Fixed-effects coefficients of a mixed-effect binomial model of the densovirus infection effect on the *Ae. albopictus* strains cannibalism. Uninfected batches of LR strain are considered as the reference level in the model (12 observations, 3 repeats).

| Fixed effects | Value | Std. Error | z-value | p-value |
|---|---|---|---|---|
| Intercept | -4.8320 | 1.3166 | -3.670 | 0.00024 |
| MTP strain | 0.2817 | 0,4381 | 0.643 | 0.52016 |
| Infection | -0.4712 | 0.5603 | -0.843 | 0.399394 |
| MTP strain : infection | 1.4526 | 0.2713 | 5.354 | 8.58e-08 |

**Table S7** - Fixed-effects coefficients of a mixed-effect binomial model of densovirus prevalence in *Ae. albopictus* and *Ae. aegypti* strains. Uninfected batches of BB strain are considered as the reference level in the model (15 observations, 3 repeats).

| Fixed effects | Value | Std. Error | z-value | p-value |
|---|---|---|---|---|
| Intercept | 1.3910 | 0.4048 | 3.437 | 0.000589 |
| LHP strain | -0.6817 | 0.5079 | -1.342 | 0.52016 |
| LR strain | -0.2314 | 0.3781 | -0.612 | 0.540473 |
| MTP strain | 0.8942 | 0.4794 | 1.865 | 0.062156 |
| SBE strain | -0.9917 | 0.4035 | -2.457 | 0.013991 |

33
